# Temperature drives variation in flying insect biomass across a German malaise trap network

**DOI:** 10.1101/2021.02.02.429363

**Authors:** Ellen A.R. Welti, Petr Zajicek, Manfred Ayasse, Tim Bornholdt, Jörn Buse, Frank Dziock, Rolf A. Engelmann, Jana Englmeier, Martin Fellendorf, Marc I. Förschler, Mark Frenzel, Ute Fricke, Cristina Ganuza, Mathias Hippke, Günter Hoenselaar, Andrea Kaus-Thiel, Klaus Mandery, Andreas Marten, Michael T. Monaghan, Carsten Morkel, Jörg Müller, Stephanie Puffpaff, Sarah Redlich, Ronny Richter, Sandra Rojas Botero, Tobias Scharnweber, Gregor Scheiffarth, Paul Schmidt Yáñez, Rhena Schumann, Sebastian Seibold, Ingolf Steffan-Dewenter, Stefan Stoll, Cynthia Tobisch, Sönke Twietmeyer, Johannes Uhler, Juliane Vogt, Dirk Weis, Wolfgang W. Weisser, Martin Wilmking, Peter Haase

## Abstract

1. Among the many concerns for biodiversity in the Anthropocene, recent reports of flying insect loss are particularly alarming, given their importance as pollinators and as a food source for many predators. Few insect monitoring programs cover large spatial scales required to provide more generalizable estimates of insect responses to global change drivers.
2. We ask how climate and surrounding habitat affect flying insect biomass and day of peak biomass using data from the first year of a new standardized distributed monitoring network at 84 locations across Germany comprising spatial gradient of land-cover types from protected to urban areas.
3. Flying insect biomass increased linearly with monthly temperature across Germany. However, the effect of temperature on flying insect biomass flipped to negative in the hot months of June and July when local temperatures most exceeded long-term averages.
4. Land-cover explained little variation in insect biomass, but biomass was lowest in forested sites. Grasslands, pastures and orchards harbored the highest insect biomass. The date of peak biomass was primarily driven by surrounding land-cover type, with grasslands especially having earlier insect biomass phenologies.
5. Standardized, large-scale monitoring is pivotal to uncover underlying processes of insect decline and to develop climate-adapted strategies to promote insect diversity. In a temperate climate region, we find that the benefits of temperature on flying insect biomass diminish in a German summer at locations where temperatures most exceeded long-term averages. These results highlighting the importance of local adaptation in climate change-driven impacts on insect communities.

## INTRODUCTION

Insects constitute much of terrestrial biodiversity and deliver essential ecosystem services such as pollination of the majority of wild plants and 75% of crop species (Losey & Vaughan, 2006; Vanbergen & Insect Pollinators Initiative, 2013). Insect biomass is a key constituent of energy flows in many food webs (Stepanian et al., 2020), a useful measure of whole insect communities (Shortall et al., 2009) and an indicator of ecosystem function (Barnes et al., 2016; Dangles et al., 2011). Climate change and anthropogenically-altered land-cover are likely drivers of insect declines, but their effects on insect biomass are still poorly characterized (Habel et al., 2019). Amidst burgeoning evidence of widespread insect declines, standardized, and large scale insect monitoring is needed to improve estimates of trends, and identify drivers (Didham et al., 2020; Wagner, 2020).

Climate change is geographically pervasive (Wilson & Fox, 2020) and may explain insect decline in natural areas (e.g. Janzen & Hallwachs, 2019; Welti, Roeder, et al., 2020). Some insect taxa are benefiting from rising temperatures, which can increase local populations (Baker et al., 2021) and range sizes (Termaat et al., 2019). However, as temperatures continue to rise and increase more rapidly, negative impacts on insect productivity are expected (Warren et al., 2018). This relationship is predicted by thermal performance theory, which hypothesizes that insect fitness, as measured by biomass or other performance indicators, will have a unimodal response to temperature (Kingsolver & Huey, 2008; Sinclair et al., 2016).

Responses of precipitation regimes to climate change vary with region, but forecasts generally suggest increased frequency of both heavy precipitation events and droughts (Myhre et al., 2019). High precipitation increases insect mortality and shortens the period of time insects are flying (Totland, 1994). Indirect effects of precipitation on flying insects mediated by plants (e.g. altering plant phenology or plant nutrition) are context-dependent but increasing rainfall in average to wet climates is often detrimental (Lawson & Rands, 2019).

Changing land-cover due to human activities is additionally a major threat to insects (Wagner, 2020), causing loss of resources and nesting locations at local scales, to fragmented habitats at larger scales (Newbold et al., 2020). Heavily human-modified landscapes come with associated pressures, such as eutrophication and pesticide use with agricultural intensification (Carvalheiro et al., 2020; Goulson et al., 2018), and urban light pollution (Owens et al., 2020), reducing both insect diversity (Fenoglio et al., 2020; Piano et al., 2020), and biomass (Macgregor et al., 2019; Svenningsen et al., 2020).

Here we ask how climate and land-cover affect flying insect biomass across the growing season and 84 locations ranging over 7° latitude during the first year of monitoring (2019) of the German Malaise Trap Program. We hypothesize **(H1)** the effect of temperature on insect biomass will **a)** be unimodal, and **b)** decline at locations where local temperatures with the greatest increase above long-term averages. We hypothesize **(H2)** that flying insect biomass will decline with increasing precipitation due to reduced flying activity. Finally, we predict **(H3)** flying insect biomass will be lower in land-cover types with larger anthropogenic impacts such as urban and agricultural areas. We additionally conducted an exploratory analysis to see if the date of peak biomass varied with climate, land-cover type, or elevation and to examine if identified significant environmental drivers of insect biomass were the result of co-variation with biomass phenology (e.g. if positive predictors resulted in capturing a phenological interval with higher biomass). The broad spatial coverage allows us to examine drivers of flying insect biomass using a macroecological gradients approach (Peters et al., 2019; Pianka, 1966).

## METHODS

### German Malaise Trap Program

The German Malaise Trap Program currently comprises 31 German Long-Term Ecological Research (LTER-D) and National Natural Landscape sites (https://www.ufz.de/lter-d/index.php?de=46285). The program was established in early 2019 to investigate long-term trends in flying insect biomass and species composition using DNA metabarcoding. In each site, one to six locations were selected and one malaise trap was installed per location. All traps measured 1.16 m^2^ on each side (Fig. S1). We examine here the 2019 biomass data retrieved from 25 of the 31 sites; the remaining sites began sampling in 2020 and are not analyzed in this study. To fill spatial gaps, we included 8 sites in Bavaria from an additional project using the same malaise trap type and measurement methods. Overall, this study includes 1039 samples from 84 locations and 33 participating sites distributed across Germany (Fig. 1; Table S1). Traps ran from early April to late October 2019 and were usually emptied every two weeks (14.03 days ± 0.06 SE; ranging 7-29 days). Some traps ran for shorter durations and several samples were lost due to animal or wind damage. By sampling across all times of day for the duration of the growing season, these data represent a wide variety of flying insect taxa across a large range of seasonal and diurnal flight periodicity.

**Figure 1.**
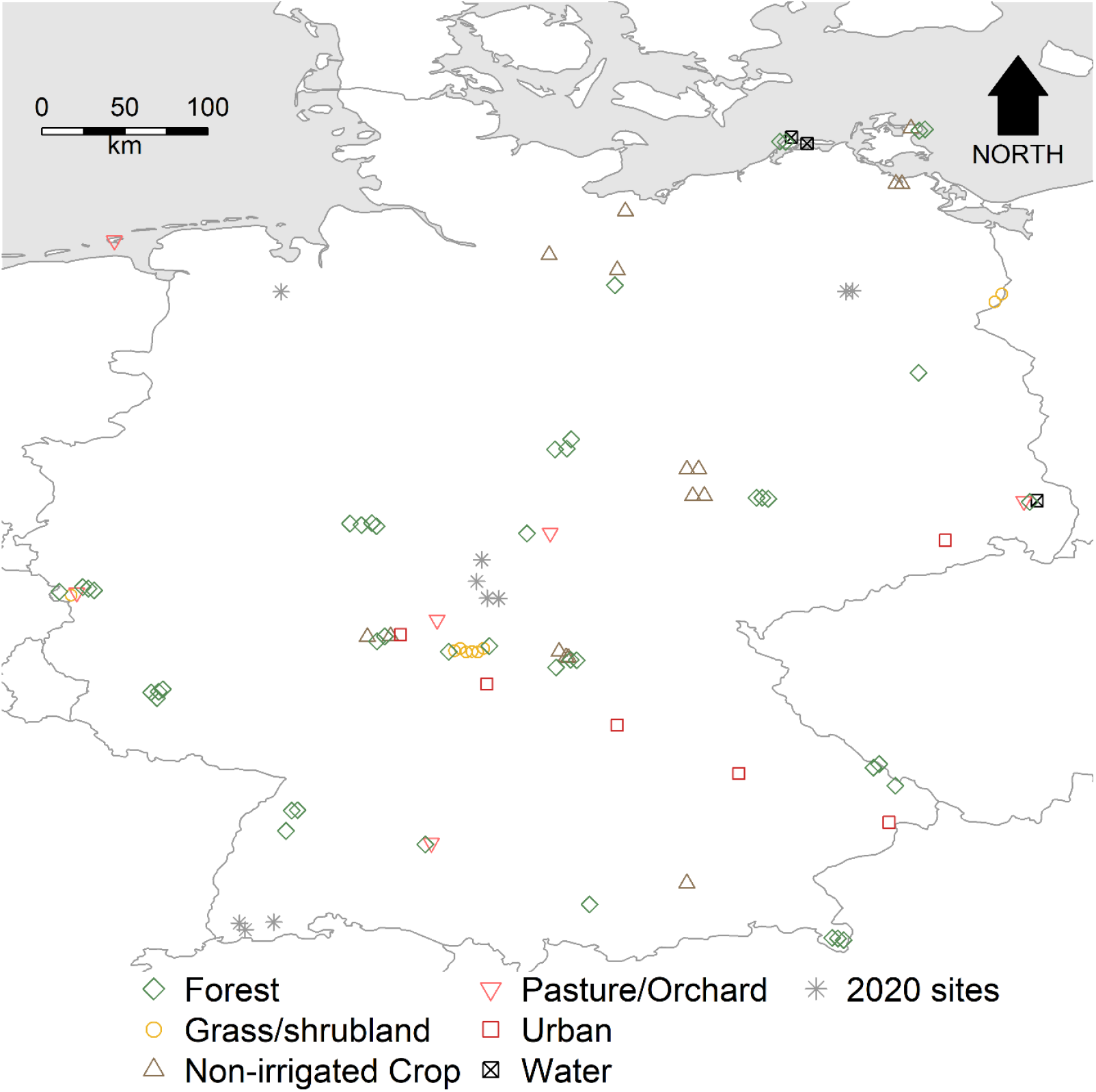
Malaise trap locations where samples were collected beginning in 2019 are identified by the dominant land-cover in the surrounding 1 km. Points coded as stars indicate trap locations at which sampling began in 2020 and are incorporated to show the full extent of the current program but are not included in the analyses. Overlapping locations were jittered longitudinally to improve visualization.

### Lab procedures

Insect biomass was wet weighed to preserve samples for future identification. Alcohol was filtered in a stainless steel sieve (0.8 mm mesh width) following the procedure in Hallmann et al. (2017), with one modification: instead of waiting until alcohol drops occurred >10 seconds apart, samples were filtered for a standard five minutes prior to weighing to the nearest 0.01g.

### Climate

Monthly means of maximum and minimum temperatures, and monthly cumulative precipitation (henceforth tmax, tmin, and precipitation) were extracted from each location from 2019 using the Terraclimate dataset (Abatzoglou et al., 2018), and from 1960-2018 using the CRU-TS 4.03 dataset (Harris et al., 2014) downscaled with WorldClim 2.1 (Fick & Hijmans, 2017). Data from both time periods (2019 and 1960-2018) were not available from either dataset alone. Both datasets have spatial resolutions of 2.5 arc minutes (∼21 km^2^) with our 84 trap locations occurring in 72 separate climate grid cells.

Tmax and tmin in 2019 were higher than 1960-2018 averages, especially during summer months (Fig. 2a) and were highly correlated (R^2^ = 0.97). Therefore, we used only tmax in our analyses. Annual precipitation was slightly lower in 2019 (784 mm ± 32 SE) relative to the 1960-2018 average (842 mm ± 32 SE), with summer months comprising the driest period, but high variation existed across months (Fig. 2b). No latitudinal temperature gradient existed across our sampling locations in 2019 (Fig. S2a) or long-term averages (Fig. S2b), likely due to a negatively correlation between elevation and latitude (Fig. S3). However, southern latitudes in 2019 experienced temperatures exceeding long-term averages to a greater degree than northern latitudes (Fig. S2c) and had higher precipitation (Fig. S2d).

**Figure 2.**
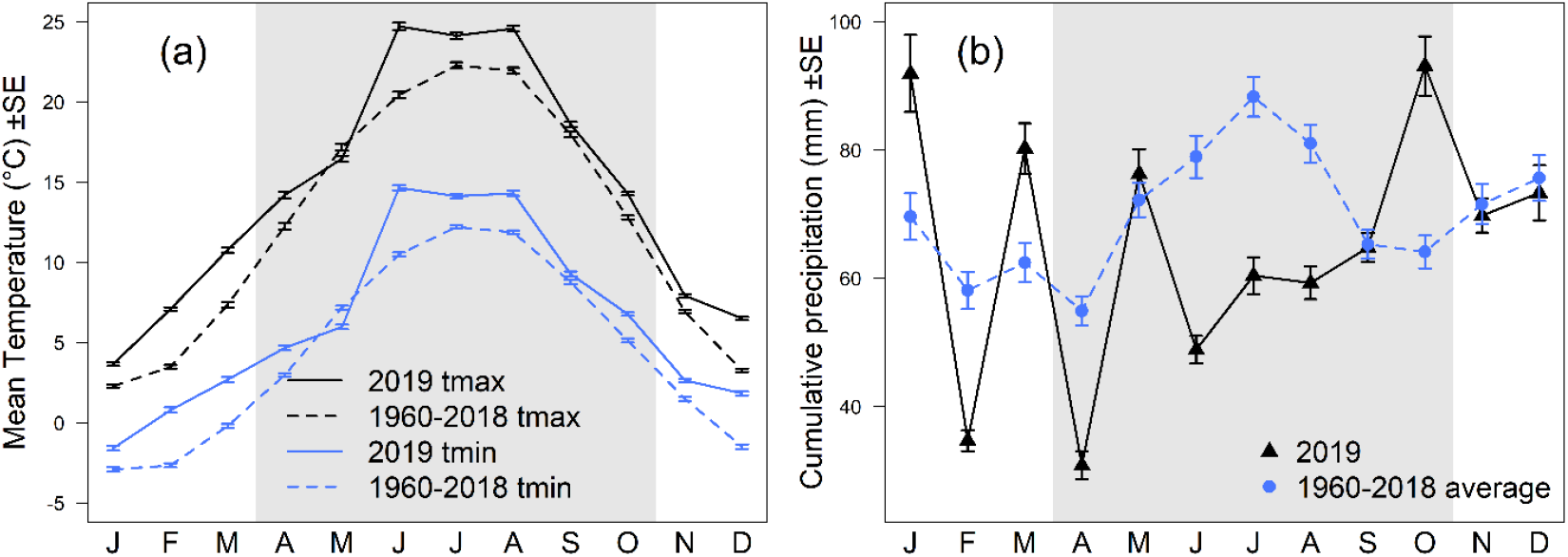
Comparison of climate at the 84 trap locations between 2019 and long-term average (1960-2018) including the average maximum monthly temperatures (tmax) and minimum monthly temperatures (tmin) in °C ± standard error (a) and cumulative monthly precipitation in mm ± standard error (b). Period of the year in which malaise trap sampling occurred is shaded in grey.

### Land-cover

Land-cover categories in a 1-km buffer around each location were extracted using the 2018 CORINE dataset (European Union, Copernicus Land Monitoring Service, 2018). Previous work suggests that at scales larger than 1-km, insects have weaker responses to land-cover buffers (Seibold et al., 2019). The 30 CORINE land-cover types were pooled into eight categories: urban (7.5% of surrounding land-cover), intensive agriculture (2.3%), non-irrigated agriculture (15.9%), pasture/orchard (12.7%), forest (44.7%), grassland/shrubland (12.1%), freshwater (3.9%), and saltwater (0.9%).

### Elevation

Elevation (m above sea level) was extracted using the Digital Terrain Model with 200-m grid widths (DGM200) from the German Federal Agency for Cartography and Geodesy (GeoBasis-DE / BKG, 2013). Elevation varied from 0 m on a barrier island in northeast Germany, to 1413 m in the German Alps.

All GIS data extraction was conducted in QGIS ver. 3.14 (QGIS.org, 2020).

### Model selection

To identify drivers of insect biomass, we used an Akaike Information Criterion corrected for small sample sizes (AIC_c_) framework (Burnham & Anderson, 2003); first building an *a priori* full model, comparing AIC_c_ of models with and without spatial autocorrelation to test for spatial non-independence, and then comparing all possible reduced models of fixed effects using the *dredge* function in the R package “MuMIn” (Bartoń, 2016). Mixed models were fit using the R package “lme4” (Bates et al., 2015). All analyses were conducted in R ver. 4.0.3 (R Core Team, 2020). To reduce variance inflation due to land-cover categories being percentages, we removed land-cover categories from the model starting with the least common until the variance inflation factor (VIF) was <10 (Montgomery et al., 2021); this removed the land-cover types of freshwater, intensive agriculture, and saltwater. VIF was calculated using the “car” package in program R (Fox & Weisberg, 2019). Initial analyses substituting the Land Use Index (LUI; Büttner, 2014) for land-cover percentages resulted in no top models containing LUI. We included the 2^nd^ degree polynomial of the sampling period to capture the season pattern of biomass. Sampling period refers to the half-month period most overlapping trap sampling days, and is numerical (e.g. first half of April = sampling period 1). Tmax and precipitation predictors correspond to the month in which the majority of sampling days occurred. Tmax was first included as a second order polynomial; however while all top models included the fixed effect of “poly(tmax,2)”, the second order polynomial term of tmax was never significant; thus we replaced this parameter with a linear “tmax” term. We initially wished to include the 2019 temperatures minus the long-term average (Δ temp) as a driver, but this variable caused inflation with sampling period and thus was excluded. Precipitation and elevation were scaled by dividing by 100.

The full model contained the response variable of sample biomass in mg/day all 84 locations and was log10(x+1) transformed to correct for a log-skewed distribution. Fixed predictors of tmax, precipitation, elevation, % cover of the five most dominant land-cover categories, the 2^nd^ degree polynomial of sampling period (poly(period,2)), and a random effect of trap location to account for repeated observations. We tested five models fitting spatial autocorrelation (exponential, Gaussian, linear, rational quadratic, and spherical correlation) and compared their AIC_c_ values with a model without a spatial correlation argument (Zuur et al., 2009). The model with the lowest AIC_c_ was the model without a spatial autocorrelation term; thus we proceeded with this model when selecting for fixed effects. Models with a ΔAIC_c_ < 2 are considered parsimonious (Burnham & Anderson, 2003) and reported.

### Temperature variation

We wished to further examine our hypothesis that the effects of temperature on flying insect biomass would decrease when local temperatures exceeded long-term averages, and examine how responses varied across sampling months. We were prohibited from including Δtemp (the deviation in monthly maximum temperatures from long-term averages) in the mixed model due to high variance inflation with sampling period. With the aim of reduce complexity due to variation in timing of sample collection across locations, and eliminate repeated sampling within a location/month, we calculated an average value of biomass (mg/day) per location and month by computing a monthly weighted average of insect biomass. Our calculation assumes traps caught the same amount of biomass each day within a sample and allocates sample biomass to each month weighed by the number of sampling days (e.g. for a trap run with 1 day in month A and 13 days in month B we assumed 1/14^th^ of the biomass was collected in month A and 13/14^ths^ was in month B). With these assumptions, the average biomass B*ij* (mg/day) of location *i* in month *j* is a weighted average of the n samples occurring in the month according to the following formula:

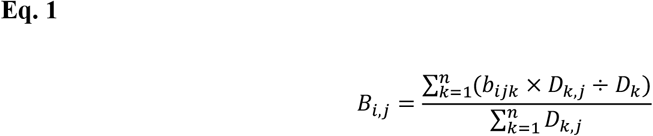

Where b_*ijk*_ = the total biomass (mg) at location *i* occurring at least partially in month *j* for a sample *k*, n= the total number of samples occurring at least partially in month *j* for location *i*, D_*k,j*_ = the number of sampling days occurring in month *j* for a given sample *k*, D_*k*_ = the total number of sampling days for a given sample *k*

For each month (April-October), we then tested for an interaction between monthly tmax and Δ temp (2019 tmax minus the long-term average tmax) for the corresponding location/month on log10 transformed B_*i,j*_. We visualized the results using the R package “effects” (Fox & Weisberg, 2019).

### Dominant land-cover categories

To visualize changes in flying insect biomass with land-cover, we plotted biomass/day over median day of sampling for locations corresponding to each dominant land-cover. Dominant land cover refers to the land cover type with the highest percentage in the 1 km buffer surrounding each location. The AIC_c_ analysis is our primary test of differences in biomass between land-cover types and uses land cover percentages rather than dominant land covers. However, we additionally used Welch’s t-tests to identify significant differences between log10(x +1) transformed B_*i,j*_ for all locations, and locations corresponding to each dominant land-cover within each month. No locations had surroundings dominated by intensive agriculture. Locations dominated by saltwater (n=1) and freshwater (n=2) were excluded due to low replication.

### Peak biomass

To calculate the day of the year of peak biomass, we fit splines on the relationship between biomass (mg/day) of each sample and the median day of the year of each sample for each location using the “smooth.spline” function in program R. We then extracted the day of the year when the maximum value of the fitted spline occurred (see Fig. S4 for an example). We excluded locations where the maximum extracted value occurred at either end of the sampling interval, assuming these sampling locations may not have captured the peak biomass date; in total we were able to calculate peak biomass date for 73 locations. We then followed the same AIC_c_ model selection procedure as was used for determining drivers of insect biomass to conduct model selection on drivers of peak biomass. The full *a priori* model was a linear regression which included the response variable of peak biomass date, and the response variables of the average monthly 2019 tmax from the beginning of the year (January) to the last main sampling month (October), the average Δtemp (2019 tmax minus long-term tmax) from January to October, the cumulative precipitation from January to October, elevation, and the % cover of the five most dominant land-cover categories. Precipitation and elevation were scaled by dividing by 100.

## RESULTS

Mean flying insect biomass averaged 2,329 ± 79 SE mg/day and varied from >10 to 17,543 mg/day. Biomass increased from 734 ± 98 SE mg/day in early April, to a peak of 5,356 ± 401 SE mg/day in late June, declining to 568 ± 111 SE mg/day in late October. AIC_c_ model comparison selected two competing top models (Table S2) with both containing tmax, percent forest cover, and sampling period, then second model additionally containing elevation as predictors of flying insect biomass (Table 1). The top model explained 43-45% of the variance in flying insect biomass without location information (marginal R^2^) and 73% of flying insect biomass was accounted for when including the random effect of location identity (conditional R^2^; Table S2).

**Table 1.** Top AIC_c_ models. AIC_c_ model selection for predictors of flying insect biomass resulted in two top models (a & b). See Table S2 for AIC_c_ parameters. Both models include the random variable of trap location. T-tests use Satterthwaite’s method. Poly(period,1) and poly(period,2) indicate the first and second order terms of the 2^nd^ degree polynomial for sampling period respectively. Other predictor variables include the percent forest in a surrounding 1 km buffer (%forest) and monthly maximum temperature (tmax). Model characteristics include estimate (Est), standard error (SE), degrees of freedom (df), t-value, and p-value (*P*).

|  | Est | SE | df | t-value | *P* |
| --- | --- | --- | --- | --- | --- |
| **a.) Model 1** |  |  |  |  |  |
| Intercept | 2.278 | 0.122 | 952 | 18.73 | < 0.001 |
| %forest | -0.319 | 0.109 | 82 | -2.93 | 0.0043 |
| poly(period,1) | -4.124 | 0.329 | 952 | -12.52 | < 0.001 |
| poly(period,2) | -4.402 | 0.707 | 952 | -6.23 | < 0.001 |
| tmax | 0.047 | 0.005 | 952 | 9.53 | < 0.001 |
| **b.) Model 2** |  |  |  |  |  |
| Intercept | 2.211 | 0.123 | 952 | 18.04 | < 0.001 |
| elevation | 0.036 | 0.013 | 81 | 2.72 | 0.008 |
| %forest | -0.487 | 0.122 | 81 | -4 | < 0.001 |
| poly(period,1) | -4.129 | 0.329 | 952 | -12.54 | < 0.001 |
| poly(period,2) | -4.288 | 0.707 | 952 | -6.07 | < 0.001 |
| tmax | 0.048 | 0.005 | 952 | 9.69 | < 0.001 |

### Climate

Flying insect biomass increased with 2019 tmax (Table 1a, Fig. S5a), and declined with increasing elevation (Table 1b, Fig. S5b). There was a significant interaction between tmax and Δtemp in the mid-season sampling months of June and July. In these two months tmax had a positive effect on flying insect biomass at locations with low Δtemps, shifting to a negative effect of tmax on flying insect biomass at locations with high Δtemps (Fig. 3; Table S3). Significant interactions between tmax and Δtemps were not found in other sampling months (Fig. 3; Table S3). The slope of the effect of temperature on flying insect biomass was steeper with lower Δ temperatures in April, August, and September, though not significantly. This pattern flipped in May and October where the slope of the effect of temperature on flying insect was steeper with higher Δ temperatures, likely due to colder temperatures in these months, though again the interaction was not significant (Fig. 3; Table S3).

**Figure 3.**
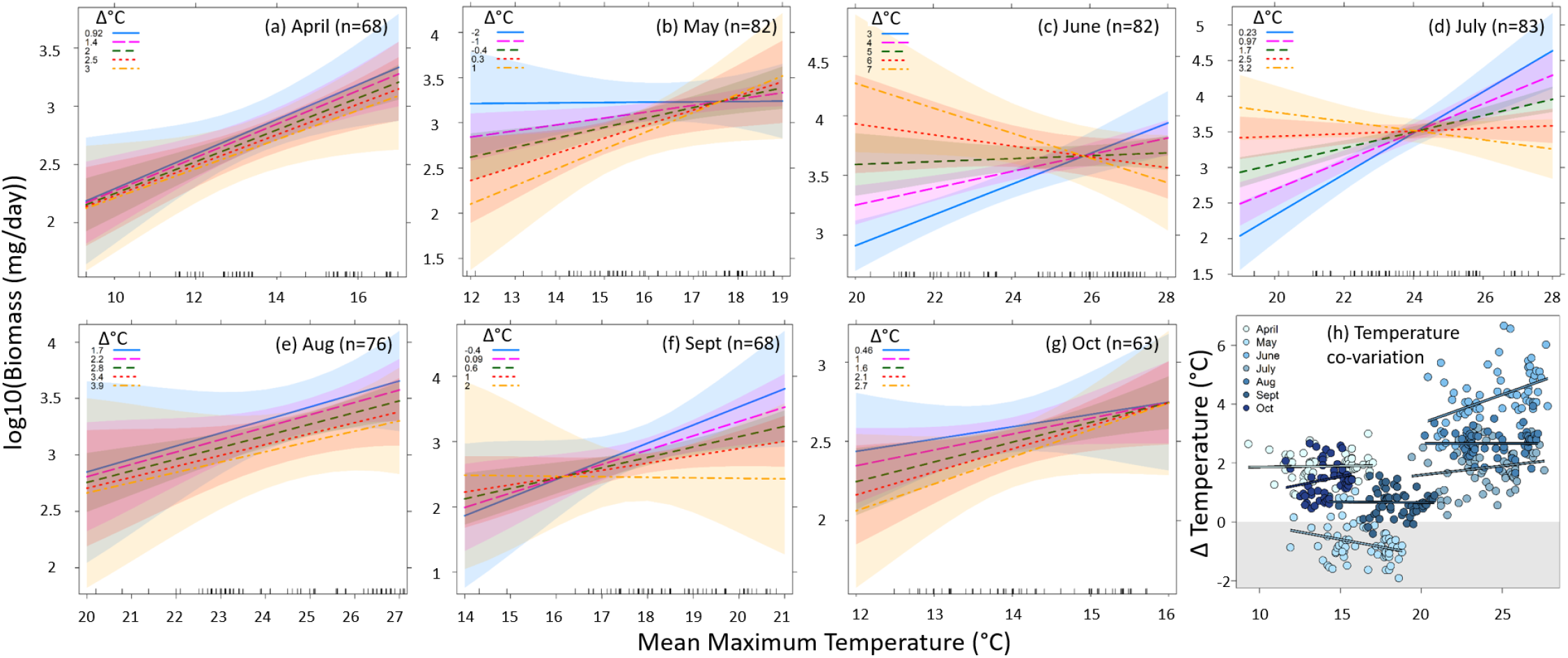
The effect of temperature on flying insect biomass was positive at the beginning of the growing season in (a) April, and (b) May regardless of Δtemp (2019 tmax minus the long-term average tmax), shifted from positive to negative with increasing Δtemp in (c) June and (d) July, and again became more positive with temperature independent of Δtemp in (e) August, (f) September, and (g) October. Number of locations with sampling (n) within each month are provided within panels a-g. While hotter months tended to have higher Δtemps, there was no consistent relationship between tmax and Δtemps within months (h). Significant interactions between tmax and Δtemp occurred in June and July; all model coefficients are provided in Table S3

### Land-cover

Flying insect biomass declined with % forest in the 1 km buffer surrounding each trap location (Table 1). No other land cover categories appeared as drivers of insect biomass. Categorizing locations by dominant land-cover suggested grassland/shrublands had the highest biomass in the mid growing season (June/July; Fig. 4c), while non-irrigated cropland supported above-average biomass at either end of the growing season (May and September; Fig. 4e). Higher biomass in urban-dominated locations (April and July-September; Fig. 4f) may be due to urban-dominated locations being in southern Germany (Fig. 1) which tended to be slightly warmer (Fig. S2).

**Figure 4.**
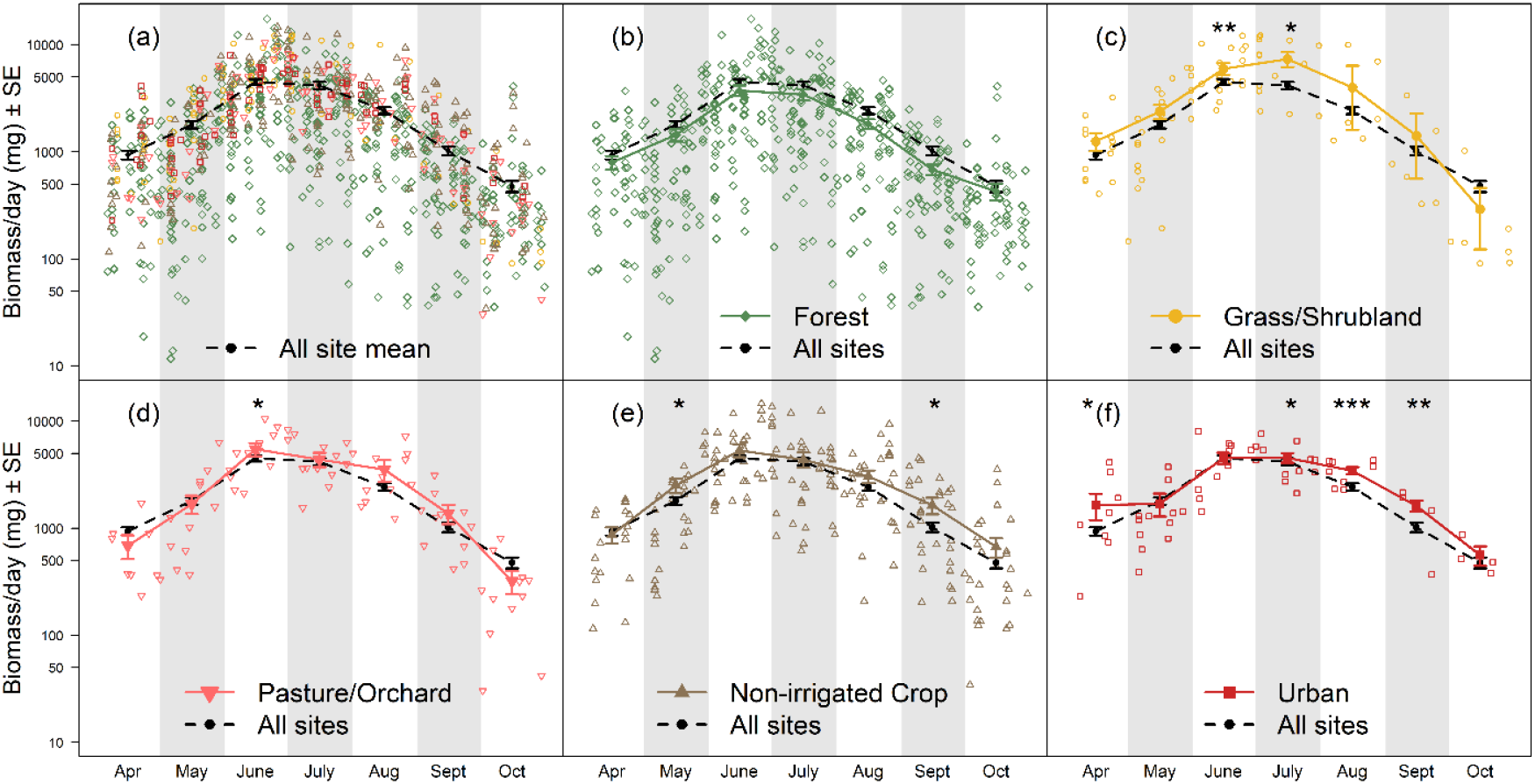
Biomass over the median sampling day across all 84 trap locations (a), and comparisons between all locations and locations with surroundings dominated by forests (b; n=44), grassland/shrubland (c; n=9), pasture/orchard (d; n=6), non-irrigated cropland (e; n=16), and urban environments (f; n=6). Point shapes and colors in panel (d) match the dominant land category following shapes and colors in panels b-f. Mean and standard error are provided for biomass within each land cover category and month. Stars indicate significant differences within each month between dominant land cover categories and all-location averages (* = 0.05 > *P* > 0.01, ** = 0.01 > *P* > 0.001, *** = *P* < 0.001).

### Peak biomass

The day of the year of peak biomass varied from 148.5 (May 28-29^th^) to 219 (Aug. 7th) across the 73 trap locations from which it was estimable (averaging 175.1 [June 24^th^] ± 1.6 days SE). Model selection resulted in 12 top models with ΔAIC_c_ < 2 (Table S4). The most consistent result was earlier peak biomass dates in locations with more surrounding grassland/shrubland. Other drivers of peak biomass date included earlier peak biomass date with increasing elevation, Δtemp, and % forest, and later peak biomass date with increasing precipitation, % pasture/orchard, and % urban surroundings. However, predictive power of top models of peak insect biomass date was low (R^2^ s ranging from 0.07-0.14; Table S4).

## DISCUSSION

In a study of 84 locations widely distributed across Germany, we found strong effects of temperature on flying insect biomass. Biomass increased linearly with temperature in contrast to the unimodal relationship predicted by the first prediction of our first hypothesis (**H1a**); however, when high, positive deviations from long-term average temperatures were combined with the hotter summer months of June and July, temperature no longer had a positive effect on flying insect biomass, in support of our second hypothesis (**H1b)**. Temperatures in June 2019 were especially hotter than long-term averages across trap locations (averaging 4.3°C ± 0.1 SE). In contrast, insect biomass only increased with temperature in May 2019, which was cold relative to the long-term averages (averaging -0.7°C ± 0.1 SE). The negative effect of high deviations from long-term temperature averages suggests insects are adapted to local temperature conditions. Rapid rises in temperature may exceed locally established tolerance limits, having negative effects on insect communities even in colder climate regions.

A decelerating benefit of temperature in locations with greater increases in temperature is consistent with previous long-term studies of insects. In a study of two surveys of ant communities across North America conducted 20 years apart, and finding that sites with the largest increases in temperature had the largest declines in colony density (Kaspari et al., 2019). Hallmann et al. (2017) found a positive effect of temperature on insect biomass; however, biomass loss over time was greatest in mid-summer, when temperatures are highest. Flying insects may be more affected by rising temperatures than non-flying insects as they cannot buffer high temperatures by burrowing in soil or plant tissue (Baudier et al., 2015; Wagner, 2020). We predict future monitoring will detect increasingly negative effects of temperature due to ongoing climate warming, as temperature begins to exceed species’ optimum temperature ranges.

Climate change predictions for Germany suggest slight increases in cumulative annual precipitation, but shifts in the timing of rainfall and drier summers (Bender et al., 2017). The 2019 growing season matched this prediction with June, July and August being much drier than the long-term average and with the wettest month of the study period being October. As insects can detect changes in barometric pressure and will stop flying if they sense storms approaching (Pellegrino et al., 2013), we predicted increased rainfall would result in reduced flight activity, reducing insect biomass. However, precipitation was not a significant predictor of flying insect biomass as predicted by **H2** potentially due to low variation in precipitation across locations.

With ∼75% of global land significantly altered by human activities (IPBES, 2019), land-cover change and land use intensification is a major contributor to insect decline (Díaz et al., 2019; Potts et al., 2010; Winfree et al., 2011). In contrast to **H3**, we did not detect negative effects of urban and agricultural land-cover on flying insect biomass. The strongest effect of surrounding land-cover was reduced insect biomass in forests. Forests may provide fewer floral resources than open fields (Jachuła et al., 2017). Alternatively, forest vegetation structure may limit insect movement through the landscape, reducing trap catch in comparison to open systems like grasslands (Cranmer et al., 2012). The absence of an effect of heavily human-impacted habitats on flying insect biomass may be due to a minority of our locations surrounded by high proportions of these land-cover types, especially intensive agriculture. Higher temperatures in urban areas may explain the above average biomass in spring and late summer/autumn, while also making insects in urban areas more vulnerable to future warming in mid-summer. Additionally, large variability exists in insect habitat quality of urban areas and agricultural land, ranging from paved expanses and areas with intensive pesticide use to urban gardens and low intensity organic farms (Bengtsson et al., 2005; Hausmann et al., 2020). While moderately impacted by human activity, non-irrigated agricultural areas, pasture land, and orchards in this study tended to support higher biomass, suggesting these land use types may provide suitable habitats for Germany’s flying insects. Alternatively, fertilization and the prevalence of monoculture on conventional farms may increase insect biomass through alleviating nutrient limitation and providing high concentrations of host plants, while not benefiting insect biodiversity (Haddad et al., 2000; Root, 1973).

While the date of peak biomass ranged from late May to early August across trap locations and varied with land-cover types, the percent variance explained by environmental drivers was low. The average temperature at trap locations was not a predictor of the date of peak biomass, suggesting the overall positive response of flying insect biomass was not driven by shifts in biomass phenology. However, top models included a weak effect of locations with higher Δ temperatures having earlier peak biomass dates. Land-cover types and temperature may also interact in their effects on flying insect biomass, though our number of trap locations is prohibitory of examining many interaction terms. Earlier peak biomass dates in grasslands and forests compared to urban areas and pasture/orchard is indicative of differences between more natural and more human-modified areas and supported by previous work finding later phenologies of butterflies (Diamond et al., 2014) and flower bloom times (Li et al., 2021) in urban areas.

### Comparison with Hallmann et al. 2017

A recent study (Hallmann et al., 2017) reported large declines in flying insect biomass from 63 German locations over 27 years. However, sampling locations greatly varied with years and the majority (58 out of 63) were clustered in central-west Germany; the sites do not have representative coverage of Germany or comprise an extensive latitudinal gradient (coverage of 2° latitude). Average insect biomass reported in Hallmann et al. (2017) varied from 9,192 mg/day in 1989 to 2,531 mg/day in 2016 (May-Sept average; no April 1989 sampling was conducted). In comparison, our traps collected a monthly average of 2,404 mg/day in May-Sept. However, Hallmann et al. (2017) used traps which were ∼51% larger (1.79 m^2^ per side) than those in this study (1.16 m^2^), suggesting higher trap catch in this study relative to the last sampling year (2016) in Hallmann et al. (2017), if trap size has an appreciable positive effect on catch. This discrepancy is most likely due to differences in sampling locations as our study cover a wider range of locations and habitats than examined in Hallmann et al. (2017), but we cannot rule out an increase in biomass of flying insects in Germany.

### Caveats

Insect biomass is a common currency ecosystem-level measure of insect productivity and is an index of energy availability for higher trophic levels. Nonetheless, from biomass alone we cannot differentiate variation in abundance, body size, species diversity, or dominance. High temperatures may reduce insect body sizes within species (Atkinson, 1994; Klockmann et al., 2017; Polidori et al., 2020) or favor smaller species (Bergmann, 1848; Daufresne et al., 2009; Merckx et al., 2018). While one long-term study of flying insects in the Netherlands found no evidence of higher rates of decline in larger species over the past two decades (Hallmann et al., 2020), larger-bodied species may have become rare earlier in the last century (Seibold et al., 2015). Climate and land-cover change may otherwise alter insect communities by favoring particular trophic levels (Welti, Kuczynski, et al., 2020), invasive (Ju et al., 2017), or pest species (Bernal & Medina, 2018). The lack of an overall unimodal relationship temperature may be a result of the coarse taxonomic (flying insects) and temporal (∼2 week) sample resolution in comparison to other studies (e.g. Kühsel & Blüthgen, 2015). Additionally, malaise traps do not collect all flying insects with larger insects like butterflies often being underrepresented. Finally, this monitoring program does not yet include multi-year coverage of flying insect trends. However, such space for time, or ecological gradients approaches have a long and fruitful history in ecology and are a useful method for providing predictions of temporal trends in the absence of time series (Peters et al., 2019; Pianka, 1966).

### Future directions: Importance of large-scale insect monitoring programs

In this first study of flying insect biomass from the German Malaise Trap Program, we find that even in a temperate climate, the positive effect of temperature on flying insect biomass diminished when combined with high positive deviations in temperature from the long-term average, and hotter mid-summer months. These interactions could not have been elucidated without growing season-long monitoring across a large number of locations and a thermal gradient. Large-scale, long-term standardized monitoring is a critical tool to disentangle potential drivers of insect decline and understand how this varies with region and taxa. Empirical studies of insect communities often lack the spatial coverage to be broadly representative across habitats (but see Jeliazkov et al., 2016; Wepprich et al., 2019). Meta-analyses have large spatial coverage, but must reckon with variable research goals and methodologies (Gurevitch & Mengersen, 2010). Spatially distributed monitoring efforts of ecological communities primarily target plants and vertebrates but not insects (Eggleton, 2020). Notable exceptions include mosquito and ground beetle monitoring by the US National Ecological Observation Network (Thorpe et al., 2016), and several regional-scale Lepidoptera monitoring programs (e.g. Dennis et al., 2019; Kühn et al., 2008; van Swaay et al., 2019). The Global Malaise Trap Program, operating since 2012 (http://biodiversitygenomics.net/projects/gmp/), and the Swedish Malaise Trap Program (operational from 2003-2006; Karlsson et al., 2020) are taxonomic treasure troves, though neither measure biomass. The German Malaise Trap Program helps to fill the gap of a distributed, standardized, and continuous monitoring program of flying insects for Germany. Malaise traps are currently being considered as a standard component of European insect biodiversity surveys, and this program provides a blueprint for a coordinated large-scale malaise trap sampling network (Haase et al., 2018). As highlighted by the recent insect decline debate (Wagner et al., 2021), comprehensive and standardized monitoring is critical to meet the challenge of unravelling insect trends and drivers in the Anthropocene.

## Acknowledgements

We thank Beatrice Kulawig, Monika Baumeister, Michael Ehrhardt, Sebastian Flinkerbusch, Michael Hinz, Reinhard Hölzel, Enno Klipp, Sebastian Keller, Linus Krämer, Gudrun Krimmer, Paula Kirschner, Beate Krischke, Johannes Lindner, Susanne Schiele, Verena Schmidt, Dragan Petrovic, Simon Potthast, Almuth Puschmann, Lena Unterbauer, Jan Weber, Roland Wollgarten, and the Auwaldstation Leipzig for assistance in the field and lab. We are grateful to the eLTER PLUS project (Grand Agreement No. 871128) for financial support to EARW and PH. This work was further supported by DFG AY12/6-4 to MA, DFG WE3081/21-4 to WWW, DFG MO2142/1-1 to MTM, and the Bavarian State Ministry of Science and the Arts.

## Supporting Information: Supplemental Tables and Figures for

Temperature drives variation in flying insect biomass across a German malaise trap network

**Table S1.**
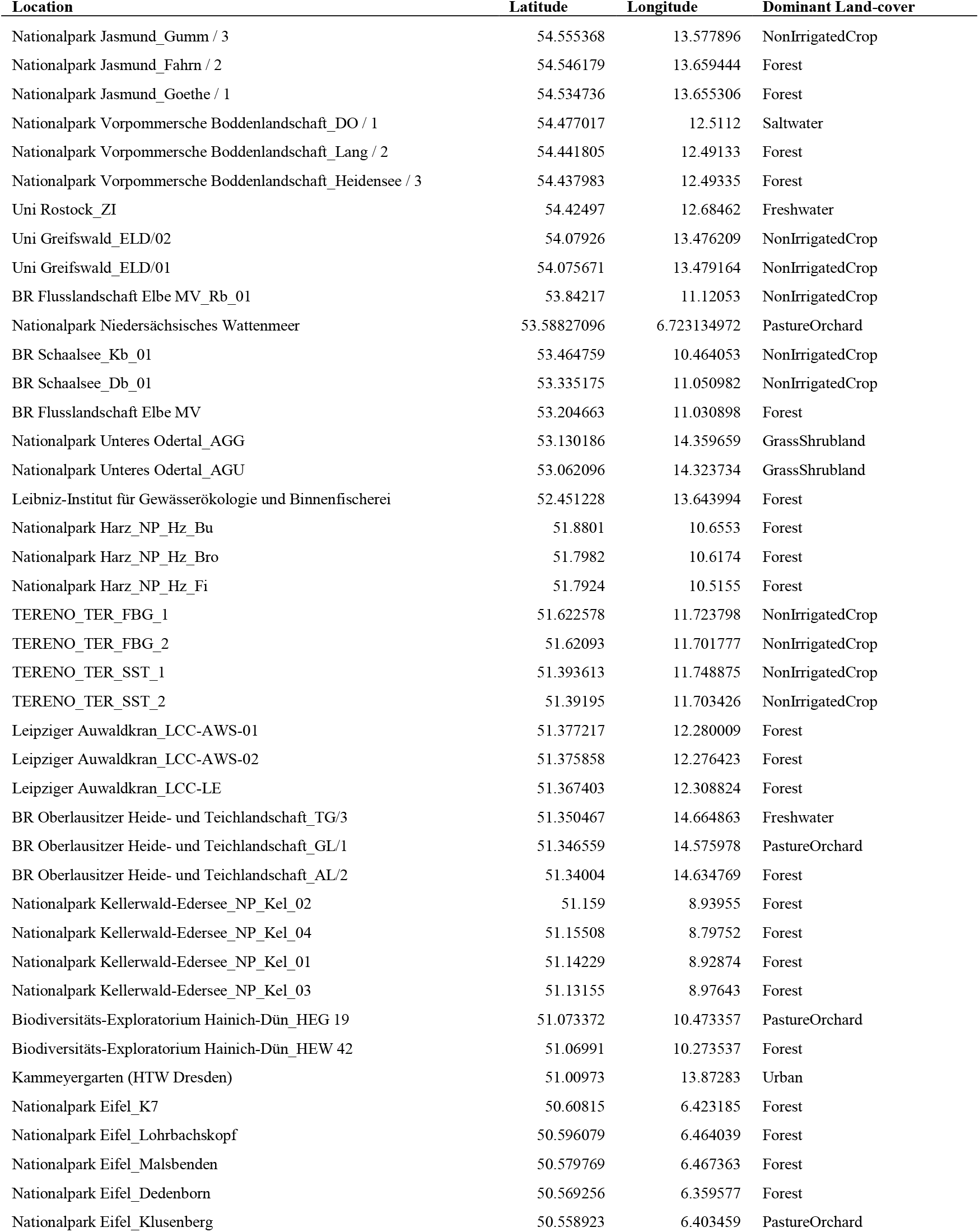
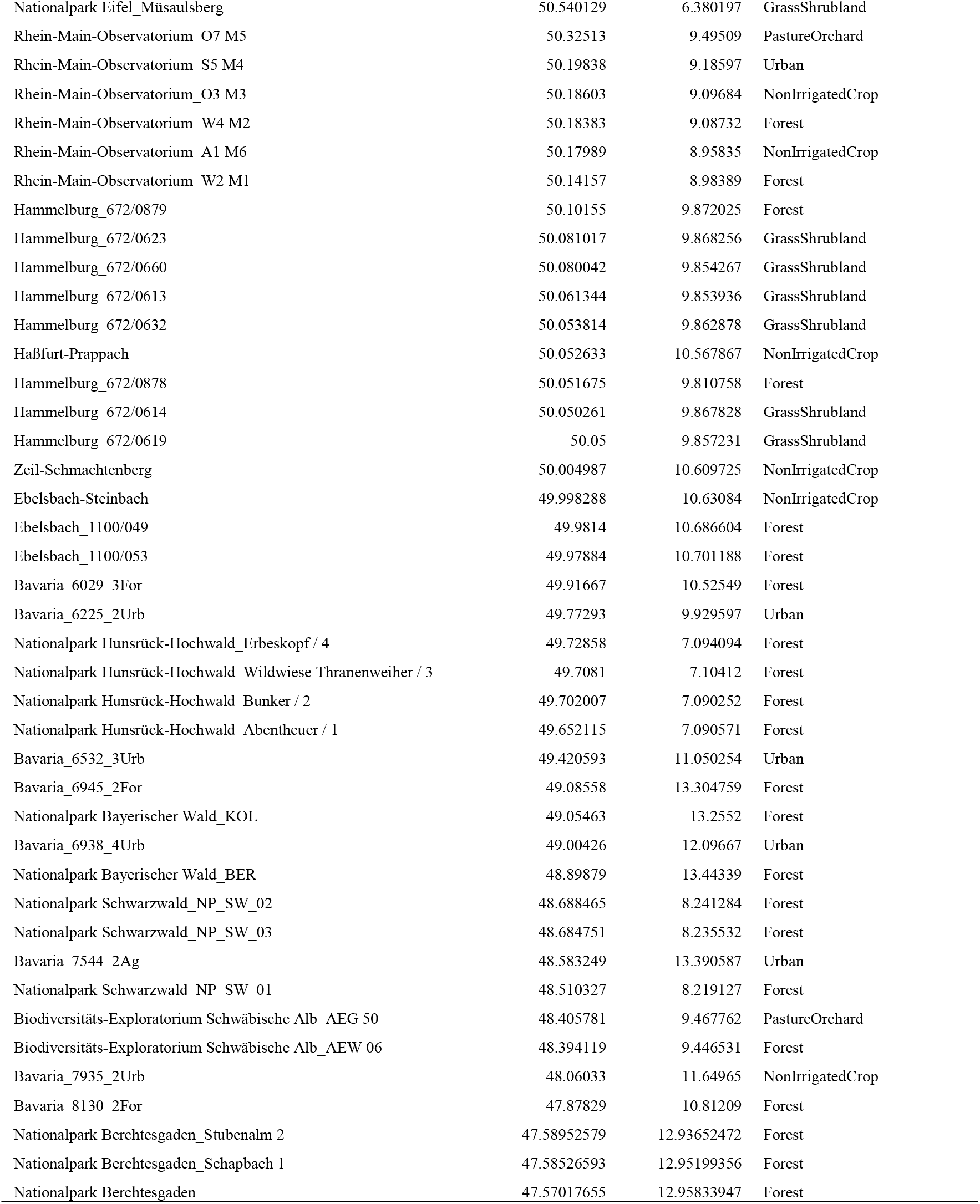
Locations of 84 malaise traps and dominant land-cover category.

**Table S2.** Top AIC models (ΔAIC_c_<2) of predictors of flying insect biomass. All models included the random variable of trap identity. Predictor variables are defined in the Methods. Marg R^2^= marginal R^2^ or the percent variance explained by the fixed effects, Cond R^2^= conditional R^2^ or the percent variance explained by the fixed effects plus the random effect of trap, df= degrees of freedom, logLik= log likelihood, and w= model weight. For summary tables of model estimates, see Table 1.

| Int | elevation | poly (period,2) | tmax | %forest | Marg $R^2$ | Cond $R^2$ | df | logLik | AICc | $\Delta$ | w |
| --- | --- | --- | --- | --- | --- | --- | --- | --- | --- | --- | --- |
| 2.28 |  | + | 0.04708 | -0.319 | 0.43 | 0.73 | 7 | -378.56 | 771.2 | 0 | 0.17 |
| 2.21 | 0.036 | + | 0.04787 | -0.487 | 0.45 | 0.73 | 8 | -378.39 | 772.9 | 1.7 | 0.073 |

**Table S3.** Model coefficients for the interaction between monthly tmax and Δtemp (Fig. 3). Models were fit for each 2019 sampling month including April (a; F_(3,64)_ = 13.2, R^2^ = 0.38, *P* < 0.001), May (b; F_(3,78)_ = 6.8, R^2^ = 0.21, *P* < 0.001), June (c; F_(3,78)_ = 14.5, R^2^ = 0.36, *P* < 0.001), July (d; F_(3,79)_ = 15.5, R^2^ = 0.37, *P* < 0.001), August (e; F_(3,72)_ = 5.3, R^2^ = 0.18, *P* = 0.002), September (f; F_(3,64)_ = 5.7, R^2^ = 0.21, *P* = 0.002), and October (g; F_(3,59)_ = 3.1, R^2^ = 0.14, *P* = 0.03).

|  | Estimate | Stand. Error | t-value | P |
| --- | --- | --- | --- | --- |
| **a) April** |  |  |  |  |
| Intercept | 0.74 | 1.42 | 0.52 | 0.61 |
| tmax | 0.16 | 0.10 | 1.53 | 0.13 |
| Δtemp | 0.07 | 0.70 | 0.11 | 0.92 |
| tmax * Δtemp | -0.01 | 0.05 | -0.22 | 0.83 |
| **b) May** |  |  |  |  |
| Intercept | 0.83 | 0.76 | 1.10 | 0.28 |
| tmax | 0.14 | 0.05 | 2.81 | 0.006 |
| Δtemp | -1.17 | 0.80 | -1.46 | 0.15 |
| tmax * Δtemp | 0.07 | 0.05 | 1.31 | 0.19 |
| **c) June** |  |  |  |  |
| Intercept | -4.19 | 1.81 | -2.31 | 0.023 |
| tmax | 0.30 | 0.08 | 4.03 | < 0.001 |
| Δtemp | 1.51 | 0.43 | 3.47 | < 0.001 |
| tmax * Δtemp | -0.06 | 0.02 | -3.28 | < 0.001 |
| **d) July** |  |  |  |  |
| Intercept | -4.11 | 1.38 | -2.99 | 0.004 |
| tmax | 0.32 | 0.06 | 5.36 | < 0.001 |
| Δtemp | 2.87 | 0.71 | 4.05 | < 0.001 |
| tmax * Δtemp | -0.12 | 0.03 | -3.94 | < 0.001 |
| **e) August** |  |  |  |  |
| Intercept | 0.32 | 4.92 | 0.06 | 0.95 |
| tmax | 0.13 | 0.20 | 0.67 | 0.50 |
| Δtemp | 0.13 | 1.75 | 0.08 | 0.94 |
| tmax * Δtemp | -0.01 | 0.07 | -0.16 | 0.88 |
| **f) September** |  |  |  |  |
| Intercept | -1.25 | 1.61 | -0.78 | 0.44 |
| tmax | 0.23 | 0.09 | 2.58 | 0.012 |
| Δtemp | 1.93 | 2.25 | 0.85 | 0.40 |
| tmax * Δtemp | -0.12 | 0.13 | -0.95 | 0.35 |
| **g) October** |  |  |  |  |
| Intercept | 1.81 | 1.75 | 1.03 | 0.31 |
| tmax | 0.06 | 0.13 | 0.46 | 0.64 |
| Δtemp | -0.66 | 1.13 | -0.59 | 0.56 |
| tmax * Δtemp | 0.04 | 0.08 | 0.51 | 0.61 |

**Table S4.**
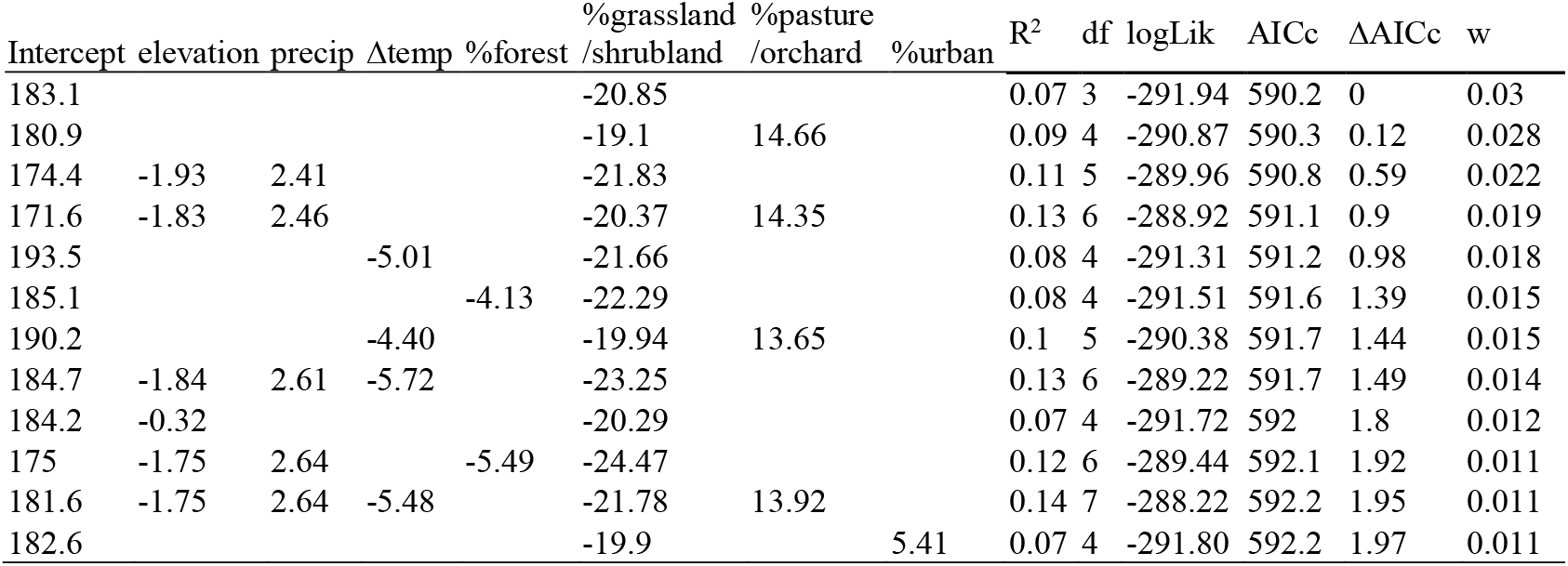
Top models (ΔAIC_c_ < 2) for day of peak flying insect biomass. Predictor variables are defined in the Methods. Df= degrees of freedom, logLik= log likelihood, and w= model weight.

**Figure S1.**
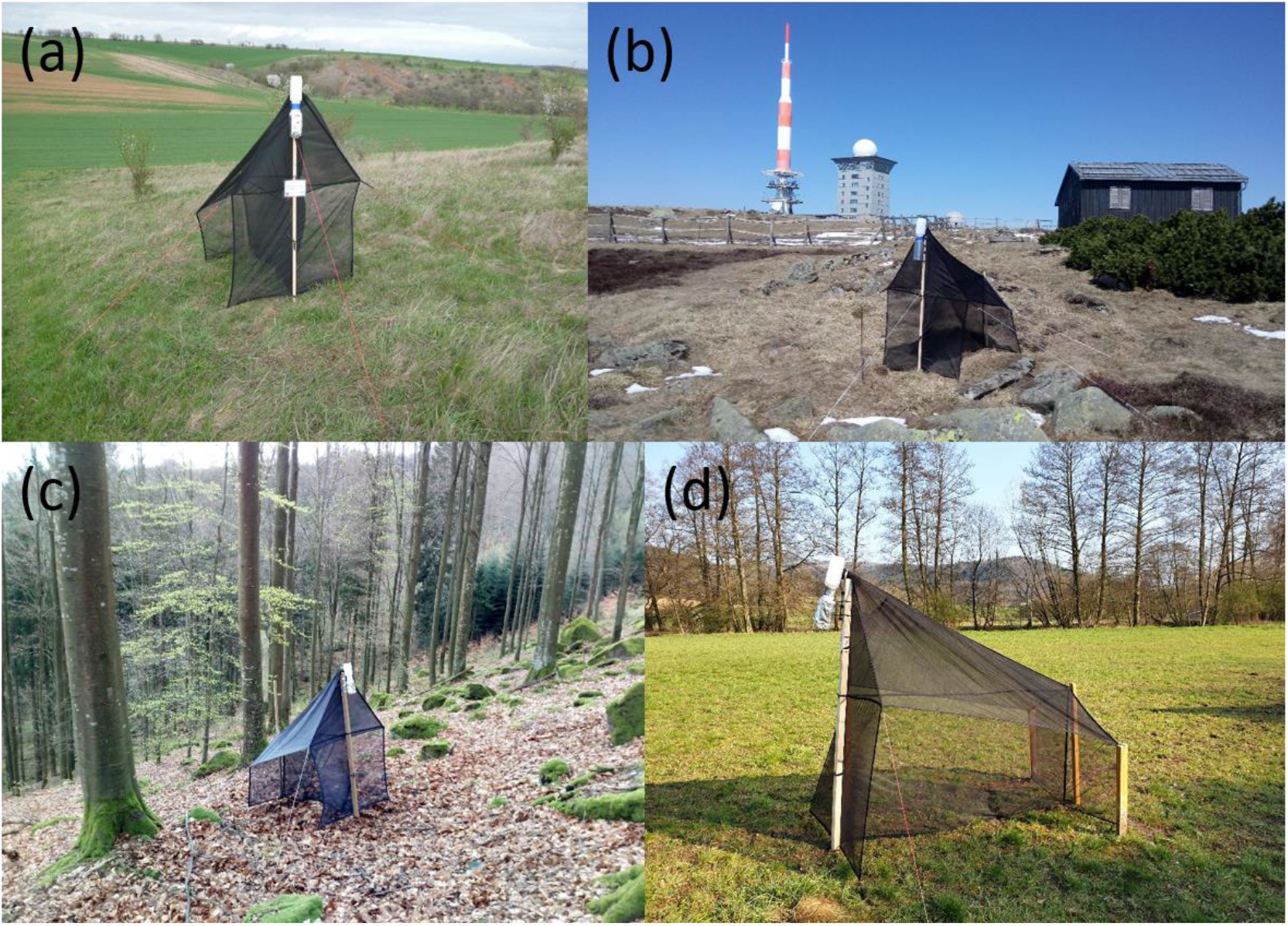
Examples of traps running in 2019 as part of the German Malaise Trap Program. Photos show traps at the LTER site Tereno-Friedeburg (a; photo credit: Mark Frenzel), at the Harz National Park (b; photo credit: Andreas Marten), at the Black Forest National Park (c; photo credit: Jörn Buse), and at the LTER site Rhine-Main-Observatory (d; photo credit: Peter Haase).

**Figure S2.**
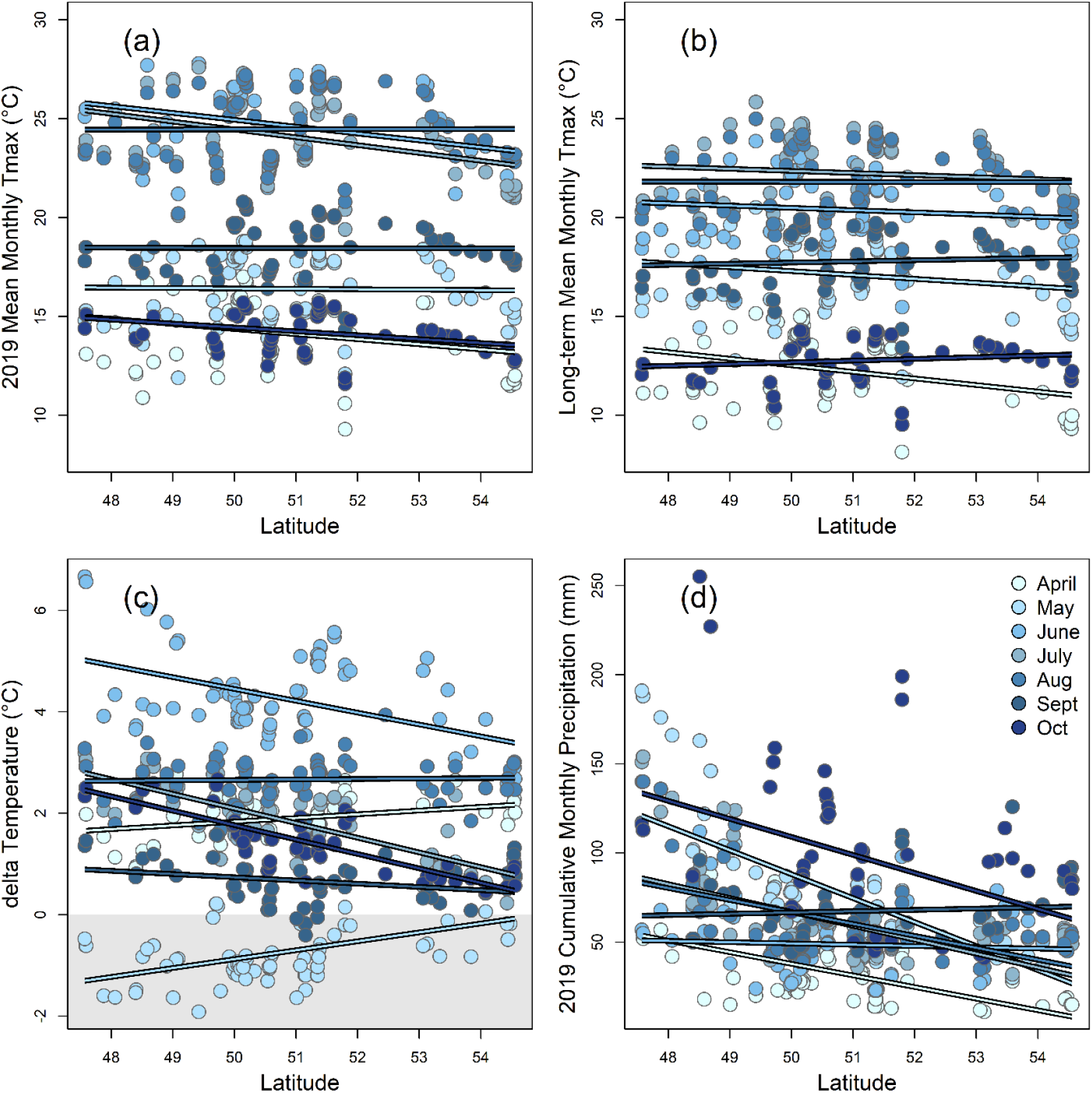
Changes with latitude across our 84 trap locations in 2019 mean monthly maximum temperature (a), the 1960-2018 long-term average monthly maximum temperature (b), the change in 2019 mean monthly maximum temperature minus the 1960-2018 long-term average (c), and 2019 cumulative monthly precipitation (d). Each point represents one month at one location, and only month/location combinations from which flying insect biomass data were collected are included. Averaging across April to October, 2019 mean monthly maximum temperature showed a weak trend to decrease with latitude (a; F_1,82_ = 2.7, R^2^ = 0.03, *P* = 0.10), while the 1960-2018 long-term average monthly maximum temperature did not vary with latitude (b; F_1,82_ = 0.6, R^2^ < 0.01, *P* = 0.44). While varying with month, the average Δ °C of 2019 maximum temperature over the long-term average decreased with latitude (c; F_1,82_ = 12.6, R^2^ = 0.13, *P* < 0.001), as did cumulative monthly precipitation (d; F_1,82_ = 26.8, R^2^ = 0.24, *P* < 0.001).

**Figure S3.**
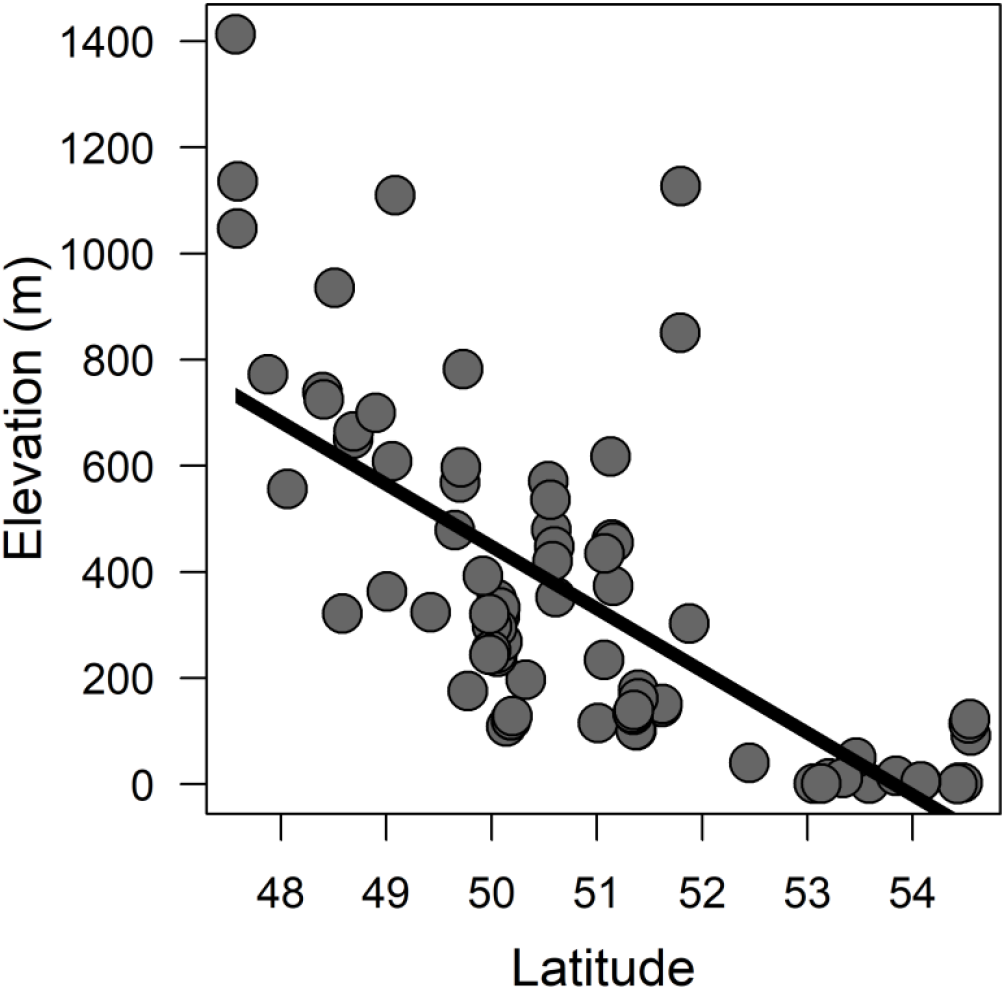
Elevation declined with increasing latitude across our 84 trap locations (F_1,82_ = 74.5, R^2^ = 0.48, *P* < 0.001).

**Figure S4.**
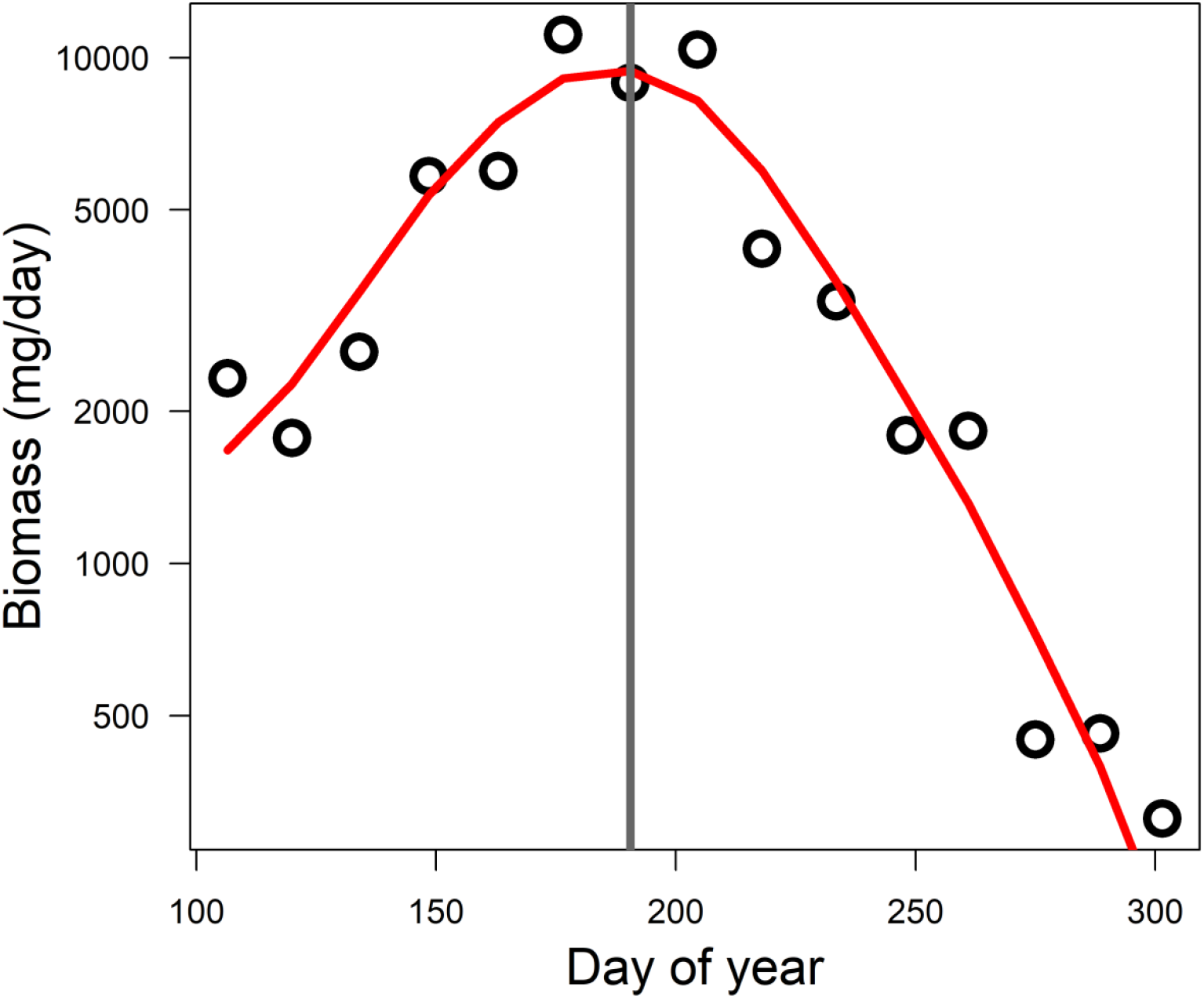
Example of determination of peak biomass day from a trap at Hunsrück-Hochwald National Park. Points represent the biomass (mg/day) collected from each sample plotted over the median day of the year of the sample. The red line is a spline fitted to these points. The grey vertical line show the maximum value of the spline, at which the day of year was extracted. At this site, 2019 peak biomass was estimated to occur at the 190.5^th^ day of the year (July 9^th^-10^th^).

**Figure S5.**
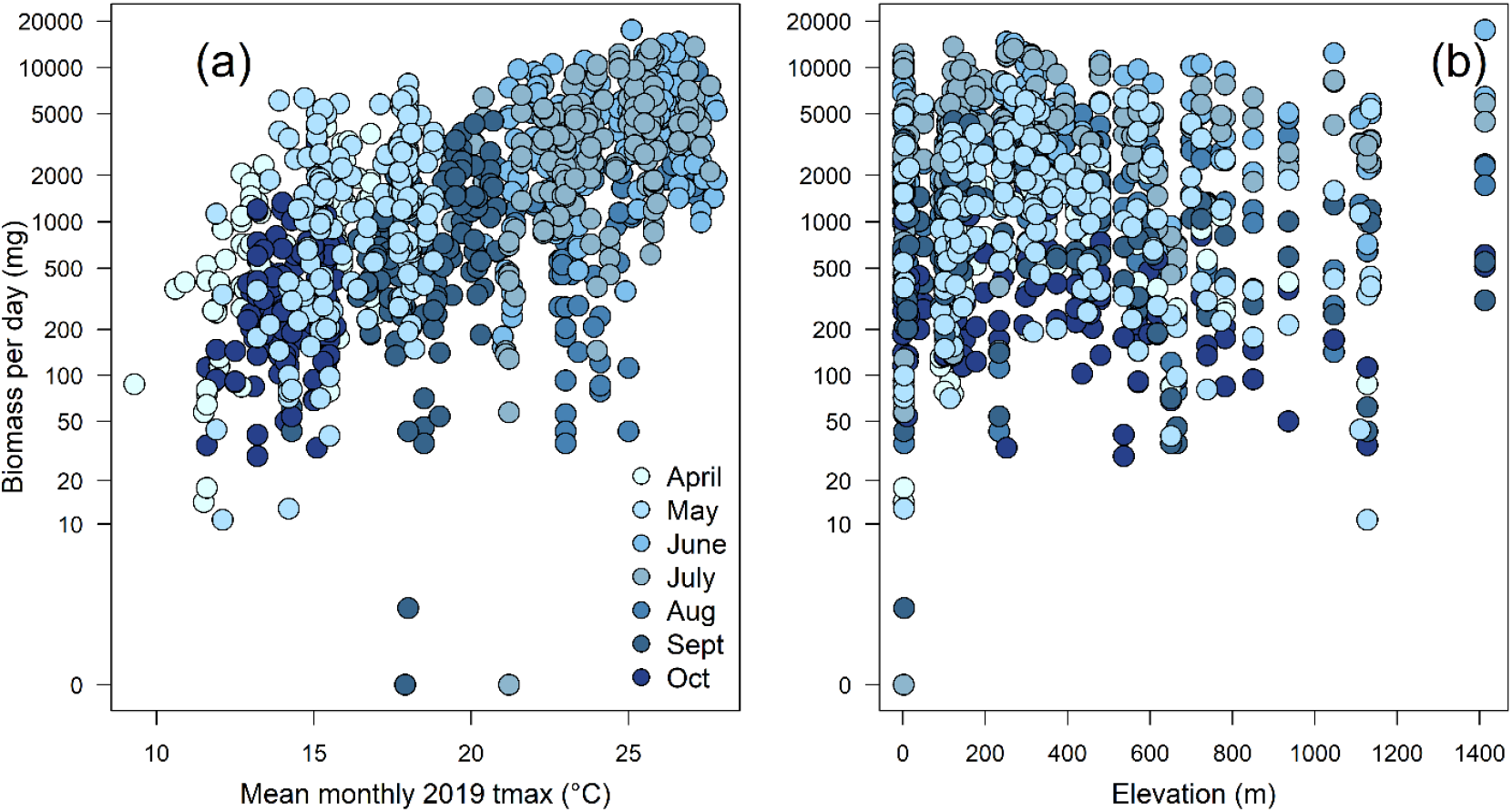
Responses of flying insect biomass to tmax and elevation. Each point represents the biomass from one sample. Across all months and site combinations, flying insect biomass increased with mean monthly 2019 maximum temperature (a), and increased weakly with elevation (b). Model estimates are provided in Table 1.

